# Epithelial restitution defect in neonatal jejunum is rescued by juvenile mucosal homogenate in a pig model of intestinal ischemic injury and repair

**DOI:** 10.1101/361352

**Authors:** Amanda L. Ziegler, Tiffany A. Pridgen, Juliana K. Mills, Liara M. Gonzalez, Laurianne Van Landeghem, Jack Odle, Anthony T. Blikslager

## Abstract

Intestinal ischemic injury results sloughing of the mucosal epithelium leading to host sepsis and death unless the mucosal barrier is rapidly restored. Neonatal necrotizing enterocolitis (NEC) and volvulus in infants is associated with intestinal ischemia, sepsis and high mortality rates. We have characterized intestinal ischemia/ repair using a highly translatable porcine model in which juvenile (6-8-week-old) pigs completely and efficiently restore barrier function by way of rapid epithelial restitution and tight junction re-assembly. In contrast, separate studies showed that younger neonatal (2-week-old) pigs exhibited less robust recovery of barrier function, which may model an important cause of high mortality rates in human infants with ischemic intestinal disease. Therefore, we aimed to further refine our repair model and characterize defects in neonatal barrier repair. Here we examine the defect in neonatal mucosal repair that we hypothesize is associated with hypomaturity of the epithelial and subepithelial compartments. Following jejunal ischemia in neonatal and juvenile pigs, injured mucosa was stripped from seromuscular layers and recovered *ex vivo* while monitoring transepithelial electrical resistance (TEER) and ^3^H-mannitol flux as measures of barrier function. While ischemia-injured juvenile mucosa restored TEER above control levels, reduced flux over the recovery period and showed 93±4.7% wound closure, neonates exhibited no change in TEER, increased flux, and a 11±23.3% increase in epithelial wound size. Scanning electron microscopy revealed enterocytes at the wound margins of neonates failed to assume the restituting phenotype seen in restituting enterocytes of juveniles. To attempt rescue of injured neonatal mucosa, neonatal experiments were repeated with the addition of exogenous prostaglandins during *ex vivo* recovery, *ex vivo* recovery with full thickness intestine, *in vivo* recovery and direct application of injured mucosal homogenate from neonates or juveniles. Neither exogenous prostaglandins, intact seromuscular intestinal layers, nor *in vivo* recovery enhanced TEER or restitution in ischemia-injured neonatal mucosa. However, *ex vivo* exogenous application of injured juvenile mucosal homogenate produced a significant increase in TEER and enhanced histological restitution to 80±4.4% epithelial coverage in injured neonatal mucosa. Thus, neonatal mucosal repair can be rescued through direct contact with the cellular and non-cellular milieu of ischemia-injured mucosa from juvenile pigs. These findings support the hypothesis that a defect in mucosal repair in neonates is due to immature repair mechanisms within the mucosal compartment. Future studies to identify and rescue specific defects in neonatal intestinal repair mechanisms will drive development of novel clinical interventions to reduce mortality in infants affected by intestinal ischemic injury.

## INTRODUCTION

The intestinal mucosa is lined by a single layer of epithelial cells, which form the principal barrier against luminal bacteria and their toxins and simultaneously facilitate selective absorption of electrolytes, water and nutrients. Intestinal ischemia leads to the breakdown of the intestinal epithelial barrier and onset of sepsis (1). Therefore, rapid and complete repair of this barrier is critical to patient survival following an ischemic event. Ischemic injury is an important contributor to intestinal mucosal disruption and inflammation in devastating neonatal diseases such as neonatal necrotizing enterocolitis (NEC), volvulus and spontaneous intestinal perforation (SIP) (2). NEC occurs in preterm infants as well as term infants with congenital heart disease and is associated with an estimated mortality rate between 20 and 30% (3, 4). Clinical interventions for neonatal intensive care unit patients with NEC currently include supportive care, attention to sepsis that results from disruption of the intestinal barrier, and ultimately intestinal resection when necessary (5). Novel treatments have focused on enhancing denovo formation of new epithelial cells, but this requires support of the patient for days following the initial injury until newly produced epithelial cells can restore the mucosal barrier (5, 6). In this subacute repair phase, remaining epithelium must immediately restitute the damaged barrier to curtail sepsis early and prevent host mortality until the regenerative phase can fully restore intestinal architecture (1). For this reason, our lab has focused on understanding the mechanisms of the subacute phase of repair, as interventions enhancing this phase will improve patient survival and hopefully reduce the need for resection in order to improve long-term quality of life.

Our lab studies mechanisms of subacute mucosal repair following ischemia using a porcine model because the pig has many fundamental anatomical, physiological, immunological and nutritional similarities to humans, and therefore provides a powerful translational model of human digestive disease, including ischemia (7–15). In juvenile (6-8-weeks-old) pigs, we have shown rapid repair of ischemia-injured mucosa involving contraction of denuded villi, epithelial cell migration across the denuded basement membrane (restitution), and re-assembly of tight junctions, resulting in swift recovery of intestinal barrier function (1, 15). Alternatively, we noted that in our studies of neonatal (2-week-old) pigs, barrier function failed to recover as efficiently following ischemic injury as compared to studies with juvenile aged pigs (16, 17). An immaturity-related defect in epithelial repair may be an important contributor to the high morbidity and mortality seen in infants affected by intestinal ischemia and barrier injury. For this reason, understanding specific rescuable defects in subacute intestinal repair mechanisms in immature patients may expedite development of novel clinical interventions to reduce mortality in infants affected by intestinal ischemic injury. Therefore, we aimed to further refine our repair model and identify rescuable defects in neonatal barrier repair. Here we describe a defect in mucosal repair in neonates that we propose is associated with insufficient pro-reparative signals from a hypomature mucosal compartment. In support of this, we demonstrate that exogenous application of injured mucosal homogenate of older animals can rescue subacute repair of injured neonatal tissue.

## METHODS

### Experimental Surgery

All procedures were approved by NC State University Institutional Animal Care and Use Committee. Two-week-old and 6-8-week-old Yorkshire cross pigs of either sex were sedated using xylazine (1.5 mg/kg) and ketamine (11 mg/kg). Anesthesia was induced with isoflurane vaporized in 100% oxygen via face mask, after which pigs were orotracheally intubated for continued delivery of isoflurane to maintain general anesthesia. Pigs were placed on a water-circulated heating pad and intravenous fluids were administered at a maintenance rate of 15 ml ⋅ kg^−1^ ⋅ h^-1^ throughout surgery. The distal jejunum was accessed via midline or paralumbar incision and 8-10 cm loops were ligated in segments and subjected to 30, 45-, 60-, and 120-minutes of ischemia via ligation of local mesenteric blood vessels with 2-0 braided silk suture, bulldog clamps, or hemostats. After ischemia, tissues were reperfused for *in vivo* recovery or loops were removed and placed in oxygenated (95% O_2_/ 5% CO_2_) Ringers solution for *ex vivo* recovery. Additional loops not subjected to ischemia were used as control tissue. At the time of loop resection, pigs were euthanized with an overdose of pentobarbital.

### Ussing chamber studies

In experiments where mucosa was stripped, the outer seromuscular layers were removed by dissection in oxygenated Ringers solution. For full thickness recovery studies, the mucosal tissue was left intact with the seromuscular layer. For *ex vivo* recovery, jejunal tissue was mounted in 1.12 cm^2^ aperture Ussing chambers. The tissues were bathed in 10 ml warmed oxygenated (95% O_2_/ 5% CO_2_) Ringer solution on the serosal and mucosal sides. Serosal Ringer solution also contained 10mM glucose while the mucosal Ringers solution was osmotically balanced with 10mM mannitol. Bathing solutions were circulated in water-jacketed reservoirs and maintained at 37°C. The spontaneous potential difference (PD) was measured using Ringer-agar bridges connected to calomel electrodes, and the PD was short-circuited through Ag-AgCl electrodes with a voltage clamp that corrected for fluid resistance. Resistance (Ω ⋅ cm^2^) was calculated from spontaneous PD and short-circuit current (I_sc_). If the spontaneous PD was between -1 and 1 mV, the tissues were current-clamped at ± 100 μA and the PD re-recorded. I_sc_ and PD were recorded every 15-minutes for 120-minutes. From these measurements, TEER is calculated. For exogenous prostaglandin experiments, 1μM 16,16-dimethylprostaglandin E_2_ was added to the basolateral chamber after the 15-minute reading for the remainder of recovery. To measure prostaglandin E_2_ production by the tissues, samples of Ringers solution were taken from the basolateral chamber to be assayed with a prostaglandin E2 metabolite ELISA kit (Cayman Chemical, catalog #514531, Ann Arbor, MI, USA). To assess barrier integrity in mucosa recovered *in vivo*, tissues were mounted on Ussing chambers in the same way but for only 60-minutes and five resistance measurements were averaged.

### Isotopic mannitol flux studies

All fluxes were conducted under short-circuit conditions (tissue clamped to 0 mV). ^3^H-mannitol (0.2 μCi/ml diluted in 10mM mannitol) was placed on the mucosal side of tissues. During *ex vivo* recovery experiments, two 60-minute fluxes from 0- to 60-minutes and from 60- to 120-minutes of the experimental recovery period by taking samples from the opposite side of that of isotope addition and counted for ^3^H β-emission in a scintillation counter. Mucosal-to-serosal fluxes (J_*ms*_) of mannitol were calculated using standard equations.(18) To assess barrier integrity in mucosa recovered *in vivo*, a single 60-minute flux was performed.

### Mucosal Homogenate and Supernatant

To obtain mucosal homogenates for *ex vivo* rescue experiments, 30-minute ischemia-injured jejunum from neonatal and juvenile pigs was harvested at the same time as tissues intended for *ex vivo* recovery. The mucosa was scraped from the jejunum using glass slides and collected into conical tubes at a ratio of 1 gram of tissue per 1ml of Ringers solution for 20-30 seconds of homogenization with a handheld tissue homogenizer (Omni International Inc., Kennesaw, GA, USA). Homogenate was then diluted to an experimental concentration of 0.2g/ml in Ringers solution, which was the highest concentration that would freely circulate through the chambers. To generate the supernatant, tubes were centrifuged at 5,400 RCF for 5-minutes and only the top layer free of solids was used. Recovery experiments began under standard protocols with Ringers solution. After the 0- and 15-minute electrical readings were recorded, the apical and/ or basolateral Ringers was drained and either the homogenates or supernatants were placed in the reservoirs, bathing the recovering neonatal mucosa from one or both sides. Glucose and mannitol were added to the homogenate or supernatant at standard concentrations.

### Light microscopy and histomorphometry

Tissues were fixed for 18 hours in 10% formalin at room temperature or 4% paraformaldehyde (PFA) in phosphate-buffered saline (PBS) at 4°C immediately following ischemic injury or after 120-minutes *ex vivo* recovery period. Formalin-fixed tissues were transferred to 70% ethanol and then paraffin-embedded, sectioned (5μm) and stained with hematoxylin and eosin for histomorphological analysis. For morphometric analysis of villus injury, the base of the villus was defined as the opening of the neck of the crypts and height of epithelialization, total height and width of villus were measured using NIH Image J^®^ Software. The surface area of the villus was calculated as previously described using the formula for the surface area of a cylinder modified by subtracting the base of the cylinder and adding a factor that accounted for the hemispherical shape of the villus tip(18). The percentage of villus epithelialization was used as an index of epithelial restitution.

### Immunofluorescence histology

PFA-fixed tissues were transferred to 10% sucrose, then 30% sucrose in PBS for 24 hours each for cryopreservation. The tissues were then embedded in optimal cutting temperature (OCT) compound and sectioned (7μm) onto positively charged glass slides for immunostaining. Slides were washed 3 times in PBS to rehydrate the tissue and remove OCT compound and were treated for antigen retrieval by in a decloaking chamber for 30 seconds at 120°C followed by 90°C for 10 seconds in a reveal decloaker solution (Biocare Medical, Concord, CA, USA). Tissues were cooled for 20-minutes at room temperature then places in PBS-0.3% Triton -100 solution for 20-minutes to permeabilize the tissues. Tissues were washed twice in PBS and incubated in blocking solution (Dako, Carpinteria, CA, USA) for 1 hour at room temperature. To mark the basement membrane, tissues were incubated in polyclonal rabbit anti-collagen IV IgG (Abcam, Cambridge, MA, USA; Catalog #ab6586) at a dilution of 1:200 in antibody diluent (Dako, Carpinteria, CA, USA) overnight at 4°C. To mark the brush border of villus enterocytes, tissues were incubated in polyclonal goat anti-villin (Santa Cruz Biotechnology, Dallas, TX, USA; catalog #sc-7672) at a dilution of 1:500 in antibody diluent (Dako, Carpinteria, CA, USA) overnight at 4°C. Intestinal fatty acid binding protein (iFABP), expressed in the cytoplasm of mature enterocytes and known to leak out during ischemic injury (19, 20), was also labeled by incubating in polyclonal goat anti-iFABP (Abcam, catalog #ab60272) at a dilution of 1:200 in antibody diluent (Dako, Carpinteria, CA, USA) overnight at 4°C. Tissues were placed in donkey anti-rabbit IgG conjugated to Alexa Fluor 568 (Invitrogen, catalog #A10042) or donkey anti-goat IgG conjugated to Alexa Fluor 488 (Invitrogen, product #A11055) at a dilution of 1:500 in antibody diluent for 1 hour at room temperature. Tissues were counterstained with nuclear stain 4′,6-Diamidino-2-Phenylindol (DAPI, Invitrogen, catalog #D1306) diluted 1:1000 in antibody diluent for 5 minutes at room temperature. Images were captured using an inverted fluorescence microscope (Olympus IX81, Tokyo, Japan) with a digital camera (ORCA-flash 4.0, Hamamatsu, Japan) using 10X objective lens with numerical aperture of 0.3 (LUC Plan FLN, Olympus, Tokyo, Japan). Specificity of primary antibodies and lack of non-specific secondary antibody binding were confirmed by secondary only negative controls.

### Scanning electron microscopy

30-minute ischemic neonate and juvenile jejunum were fixed at 0-, 30- and 120-minutes during the experimental recovery period in separate experiments. Mucosa was rinsed briefly with PBS to remove surface debris followed by immersion fixation in 2% paraformaldehyde/2.5% glutaraldehyde/0.15M sodium phosphate buffer, pH 7.4. Specimens were stored in the fixative overnight to several days at 4°C before processing for SEM (Microscopy Services Laboratory, Dept. of Pathology and Laboratory Medicine, UNC, Chapel Hill, NC, USA). After three washes with 0.15M sodium phosphate buffer (PBS), pH 7.4, the samples were post-fixed in 1% osmium tetroxide in PBS for 1-hour and washed in deionized water. The samples were dehydrated in ethanol (30%, 50%, 75%, 100%, 100%), transferred to a Samdri-795 critical point dryer and dried using carbon dioxide as the transitional solvent (Tousimis Research Corporation, Rockville, MD). Tissues were mounted on aluminum planchets using silver paste and coated with 15nm of gold-palladium alloy (60Au:40Pd, Hummer X Sputter Coater, Anatech USA, Union City, CA). Images were taken using a Zeiss Supra 25 FESEM operating at 5kV, using the SE2 detector, 30μm aperture, and a working distance of 10 to 12mm (Carl Zeiss Microscopy, LLC, Peabody, MA).

### Statistical Analysis

All data was analyzed using SigmaPlot^®^ (Systat^®^; San Jose, California, USA) and Prism^®^ (GraphPad^®^; La Jolla, California, USA) statistical software. Data were reported as means ± SE for a given number (*n*) of animals for each experiment. Results were analyzed by student’s t-test, or two- or three-way ANOVA on repeated measures. For analyses where significance was detected by ANOVA, Tukey’s test or Sidak’s test was utilized for post hoc pairwise multiple comparisons. The α-level for statistical significance was set at P<0.05. Where a significant time by treatment interaction was found, a one-way ANOVA was performed to identify individual treatment effects.

## RESULTS

### Ischemia induces time-dependent injury that is similar in neonatal and juvenile pigs

In order to determine the effect of postnatal age on level of injury following intestinal ischemia, neonatal and juvenile aged animals were subjected to jejunal ischemia for varying durations and intestinal barrier integrity was studied. In both neonatal and juvenile jejunum, 30-, 45-, 60-, and 120-minutes of ischemia induced villus contraction (Fig. 1A) and time-dependent epithelial injury (Fig. 1B). As expected, increasing durations of ischemia led to increasing injury (decreasing epithelial coverage), and there was a significant reduction in villus height after recovery in both neonates and juveniles (Fig. 1A). However, there was no significant difference in injury between neonates and juveniles within each injury duration when measured as percent epithelial coverage (fig. 1B).

**Figure 1.**
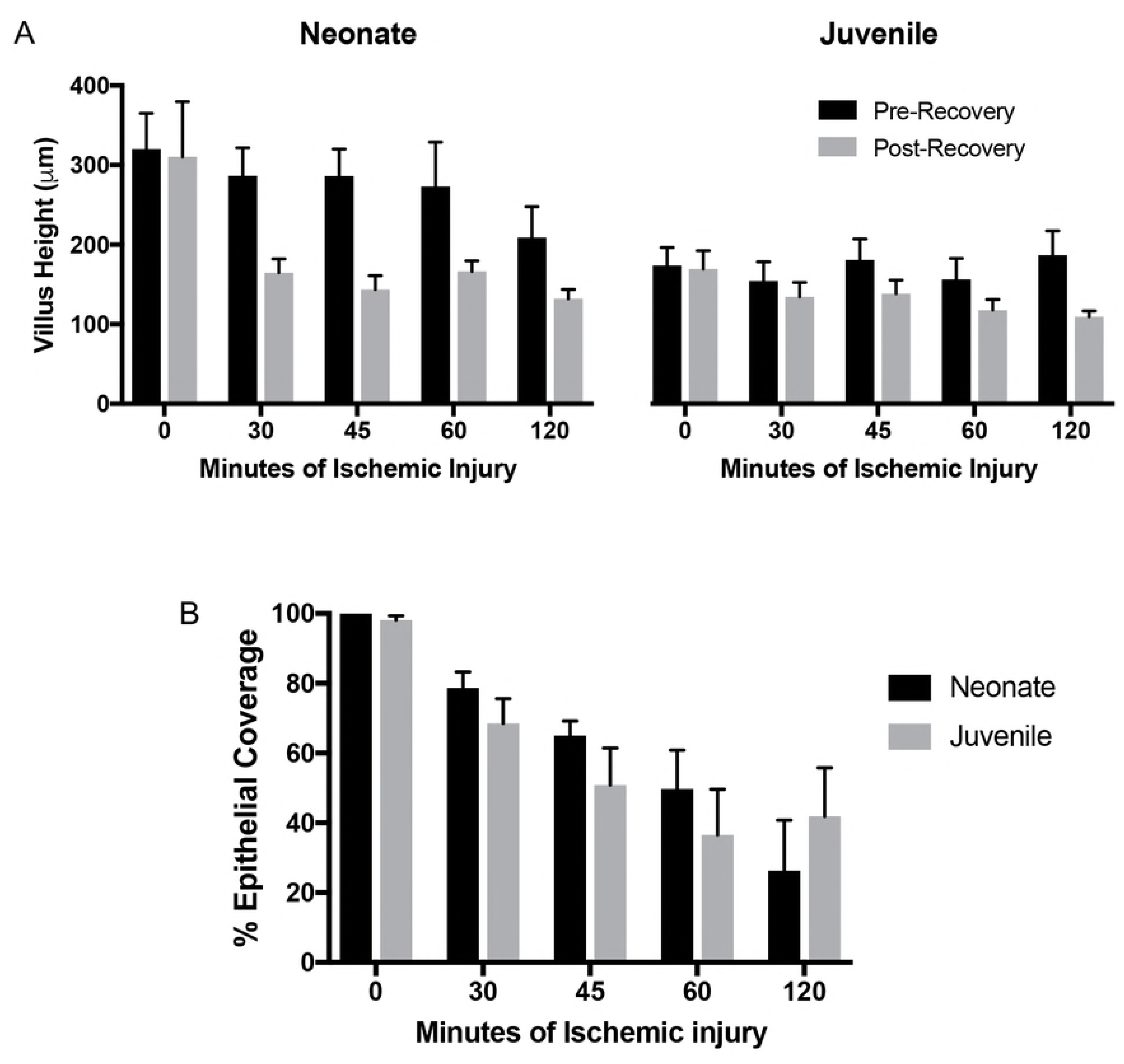
Villi show similar morphometric changes following ischemic injury in neonates and juveniles. (A) Villus height decreases following injury and the villi shorten further following 120-minutes *ex vivo* recovery in both neonates and juveniles (Significant effect of recovery and injury duration on villus height by two-way ANOVA in neonates, P<0.05; significant effect of recovery on villus height by two-way ANOVA in juveniles, P<0.05). (B) Histomorphometry quantified a similar decrease in epithelialization (increase in injury) with increasing durations of ischemia in both age groups (significant effect of ischemia on epithelialization by two-way ANOVA, P<0.0001, no significant differences between age groups at each duration of injury following Sidak’s multiple comparisons test).

To further characterize the degree of mucosal injury induced by 30-minutes ischemia in the two age groups, fixed-frozen sections were immunolabeled for enterocyte and extracellular matrix markers. Villin, which shows a strong immunoreactivity in the brush border of mature enterocytes on the villi, and collagen IV, the primary matrix component of the basement membrane, were probed in control and ischemic mucosa from both neonates and juveniles. Both age groups showed similar disruption of the enterocyte brush border due to epithelial sloughing restricted mainly to the villus tips (fig. 2A, green). A continuous lining of collagen IV at the denuded surfaces of the villus tips indicates that there was no loss of subepithelial tissue due fracturing of the villi during ischemic injury in either the neonate or juvenile mucosa (fig. 2A, red). As a marker of mature enterocyte cytoplasm as well as a functional marker of intestinal ischemic injury(19, 20), iFABP was also probed. The iFABP signal was restricted to the cytoplasm of enterocytes and was more highly expressed towards the villus tips in both age groups. Wound-adjacent enterocytes expressed iFABP in both neonates and juveniles (fig. 2B).

**Figure 2.**
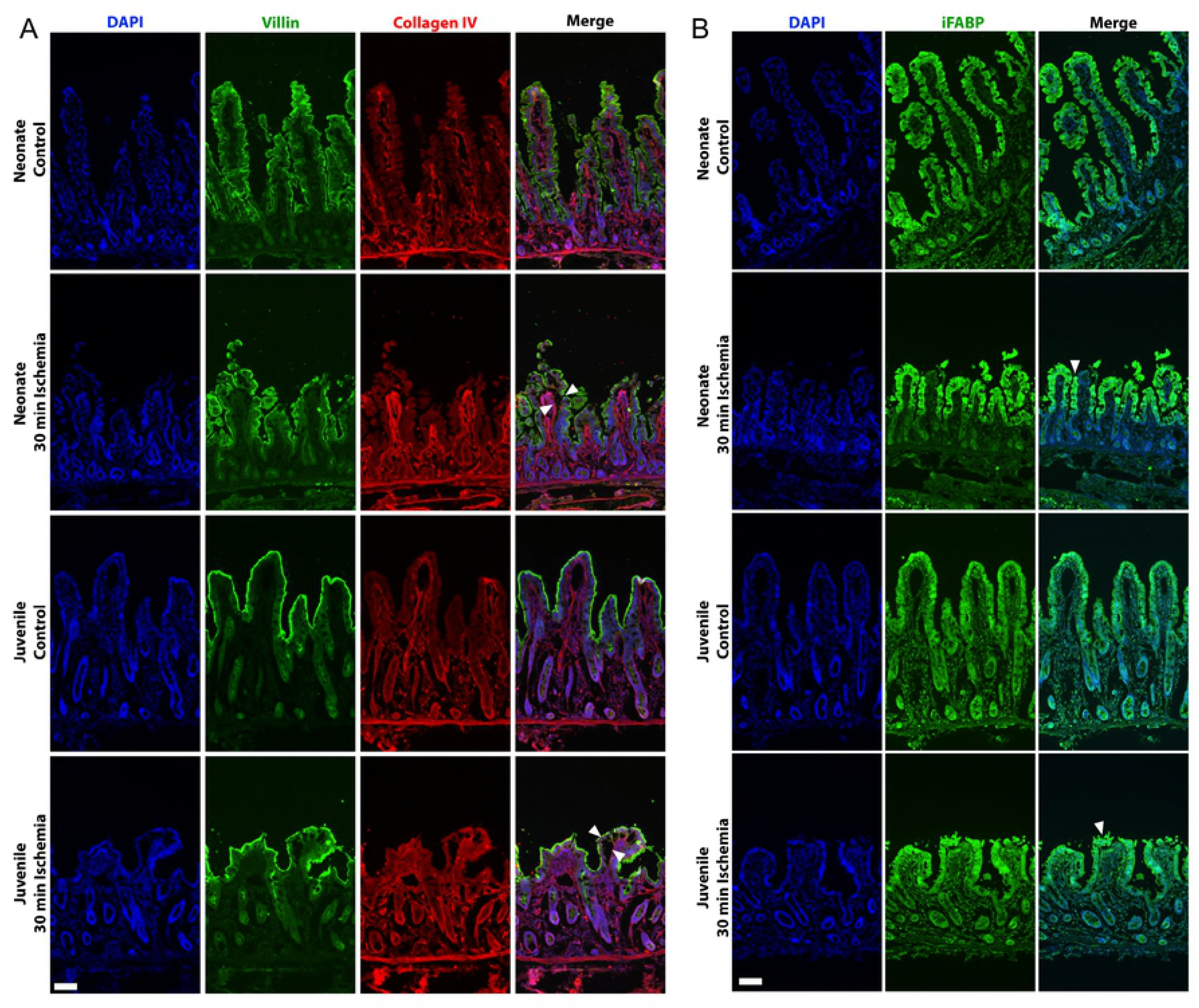
Ischemia induced similar epithelial injury in both age groups. (A) Fixed-frozen sections of neonate (top rows) and juvenile (bottom rows) jejunal mucosa was probed for Collagen IV (red), a component of the basement membrane, and villin (green), as a marker of the brush border of mature villus enterocytes. Note that ischemia-induced enterocyte loss is restricted to mainly the villus tips (white arrows), and that the basement membrane is intact in the injured mucosa from both age groups (representative fields from n=3 per group, Scale bars 100 μm). (B) iFABP (green) was restricted to the cytoplasm of enterocytes with increased expression toward the villus tips in both age groups. Note that wound-adjacent enterocytes (white arrows) express iFABP in injured villi from both neonate and juveniles (representative fields from n=3 per group, Scale bars 100 μm)

### Failure of *ex vivo* barrier repair in neonates is due to a defect in restitution

We next wanted to determine if there was a difference in the ability of neonatal and juvenile tissues to repair, as suggested by our previous studies (16, 17). Mucosal tissues were stripped from the seromuscular layers in preparation for *ex vivo* recovery in Ussing chambers. At the beginning of *ex vivo* recovery, all injured tissues of both age groups demonstrated reduced TEER as compared to age-matched controls (Fig. 3A). In juvenile tissues, TEER increased to the level of control tissues in both the 45- and 60-minute-injured groups, and the 30-minute-injured group exceeded the control levels during the 120-minute *ex vivo* recovery period. Tissues injured by 120-minutes ischemia increased but did not meet control TEER levels during recovery (Fig. 3A, right panel). In neonatal tissues, there was no change in TEER throughout the entire *ex vivo* recovery period regardless of duration of ischemic injury (Fig. 3A, left panel). The most notable differences between neonates and juvenile animals were seen when comparing the 30-minute injured jejunum, where there was a significant difference in TEER of juvenile tissues after 90-minutes of recovery versus neonatal tissues (Fig. 3B). This was accompanied by a decrease in ^3^H-mannitol flux from the first hour to the second hour of the recovery period in juveniles as compared to no change in neonates (Fig. 3C). Histology revealed a remaining defect in the intestinal epithelium at the villus tips following recovery of injured neonatal jejunum as compared to a ‘macroscopically sealed’ layer of newly restituted epithelium at the villus tips in juveniles (Fig. 3D). In 30-minute ischemia injured tissues, histomorphometry quantified 93% wound closure in juveniles as compared to an 11% increase in wound size in neonates following recovery (Fig. 3E). Due to the notable differences in barrier recovery in juveniles versus neonates following 30-minutes ischemia, all studies that follow utilized 30-minutes of jejunal ischemia.

**Figure 3.**
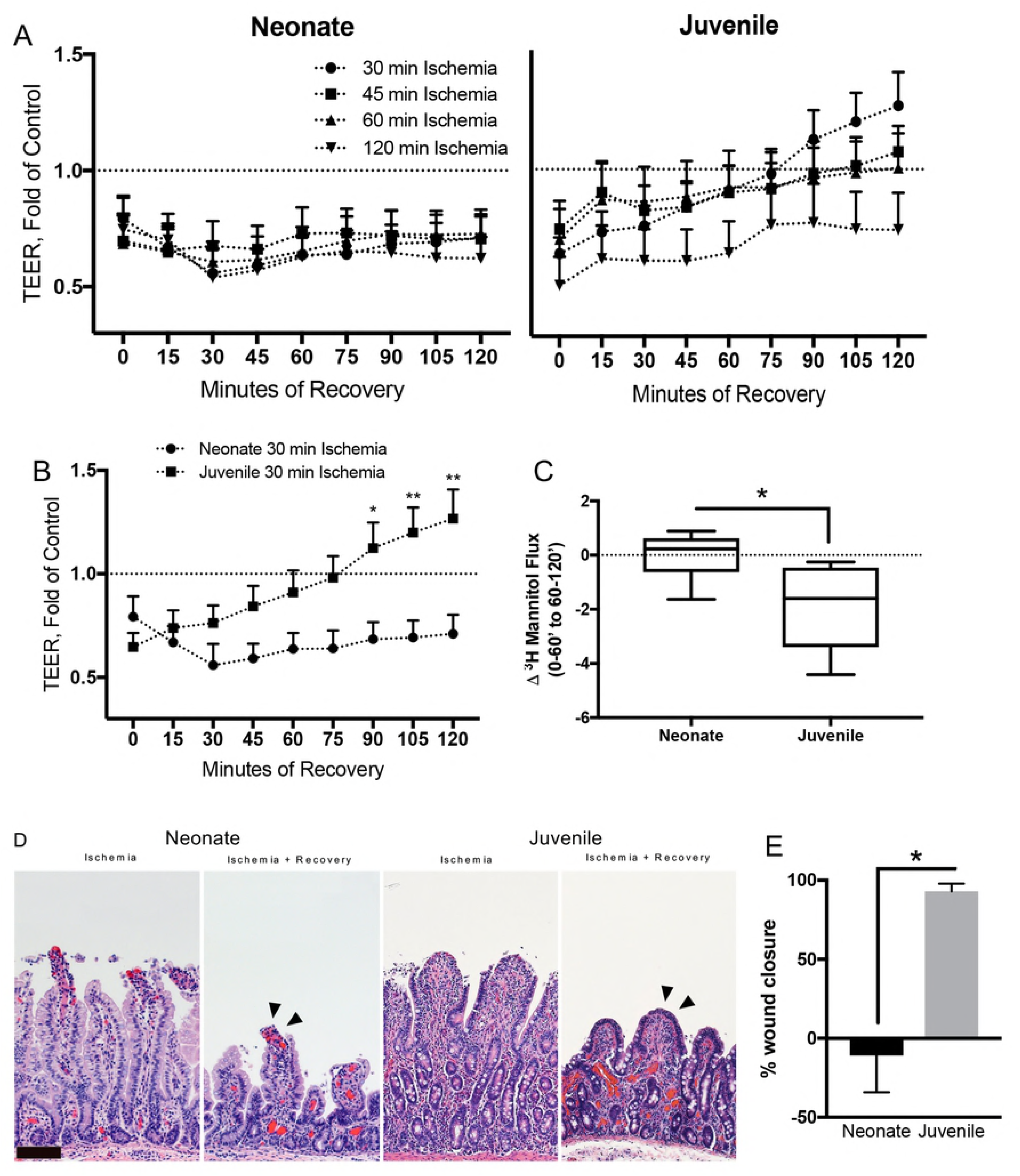
Neonates fail to recover barrier function ex vivo following ischemia due to failure of restitution. (A) TEER over 120-minutes ex vivo recovery in Ussing chambers. Note the return of TEER beyond control values in the 30-, 45-, and 60-minute ischemia-injured juvenile mucosa within 120-minutes of *ex vivo* recovery, while the all ischemia-injured neonatal tissues fail to recover TEER within 120-minutes (n=5-17, significant interaction between recovery and age on three-way repeated measures ANOVA, P≤0.001). (B) TEER of 30-min ischemia-injured tissues versus controls over 120-minutes *ex vivo* recovery (two-way ANOVA with Sidak’s multiple comparisons test, ^∗^P≤0.05, ^∗∗^P≤0.01). (C) Change in ^3^H-mannitol flux relative to controls in 30-min ischemia-injured tissues over the *ex vivo* recovery period. Note no change in small molecular flux in neonates whereas juvenile tissue flux decreases from the beginning to the end of the recovery period (n=6, Student’s t-test, ^∗^P≤0.01). (D) Representative histology shows the remaining defect in the 30-minute ischemia-injured mucosal epithelium at the villus tips in neonates after recovery as compared to the restituted epithelium (arrowheads) of the juvenile villi (scale bar 100 μm). (E) Histomorphometry quantified a 93±4.7% wound closure in juveniles as compared to a 11±23.3% increase in wound size in neonates (n=6-18, ^∗^P≤0.05, student’s t-test).

### Neonatal wound-associated epithelial cells fail to assume a migratory phenotype

Based on the lack of restitution noted, we next wanted to assess the nature of the wound-adjacent enterocytes more closely to determine whether these cells were undergoing phenotypic changes associated with epithelial restitution. To do this, ischemia-injured villus tips undergoing *ex vivo* recovery were visualized by scanning electron microscopy. Initially after injury, both neonate and juvenile mucosa exhibited damaged and sloughing enterocytes at the villus tips. Contrasting to the juvenile shorter villi, the longer neonatal villi formed concentric folds in the intact epithelium indicative of the contraction of the villus core beneath the surface (Fig. 4A). In juvenile tissues, enterocytes at the defect margins assumed a migratory phenotype characterized by the depolarization/ flattening of the cells, loss of microvilli, and extension of lamellipodia into the wound bed (Fig. 4B, right panels). Interestingly, in neonatal tissues, the cells assumed an atypical spherical shape (Fig. 3B, left panels) which to our knowledge has not been reported before. These spherical cells did not appear to assist wound closure, but rather remained at the edges of the wound. Indeed, in contrast to juvenile tissues, enterocytes beside the wound bed retained a polarized phenotype with no evidence of assuming any of the features of restitution. More specifically, they did not flatten, retained microvilli, and lacked the lamellipodial extensions indicative of cell crawling.

**Figure 4.**
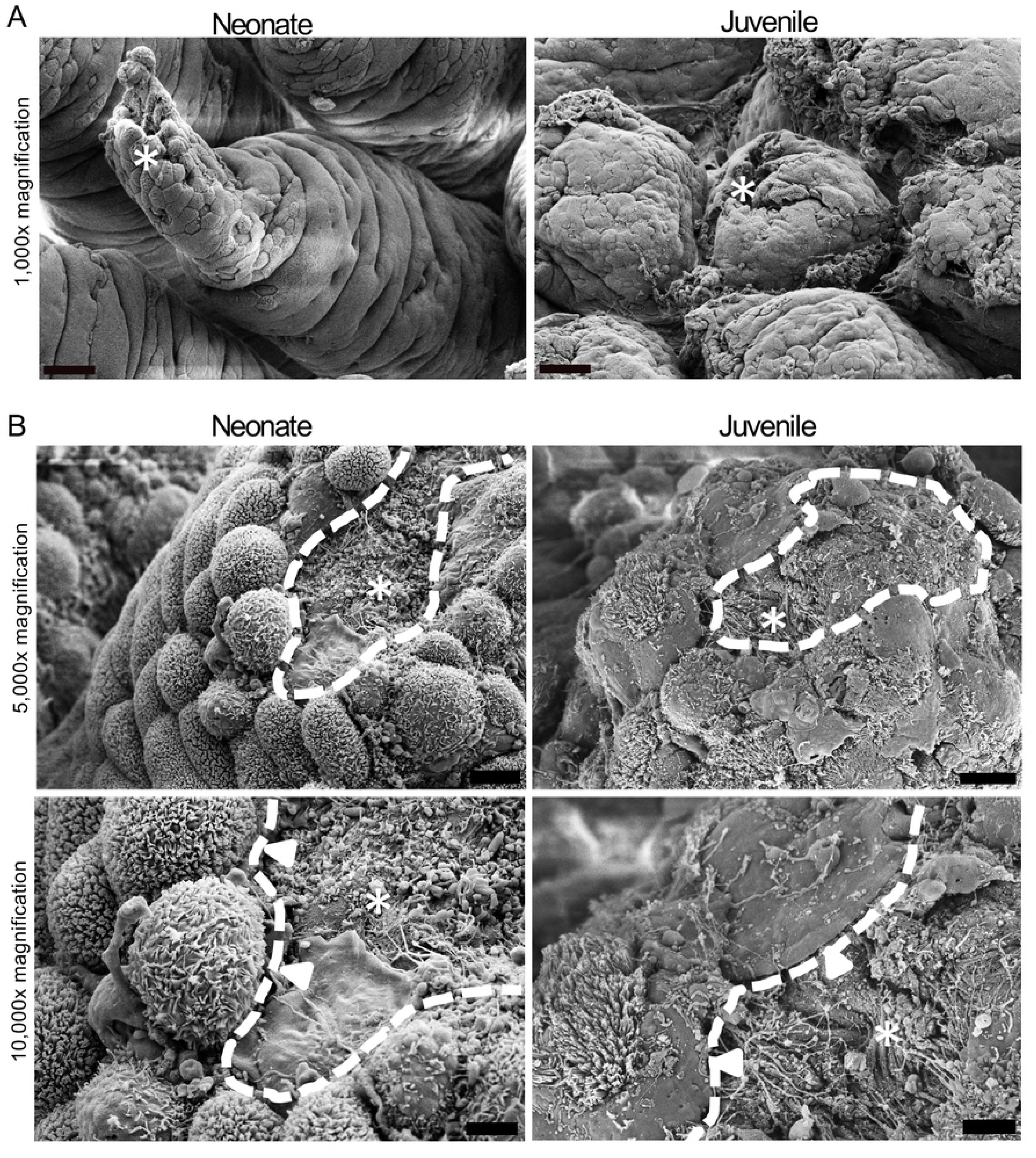
Scanning electron microscopy shows neonatal wound-associated enterocytes fail to assume migratory phenotype seen in juveniles. (A) Neonate and juvenile villi following 30-minutes ischemia. Note the sloughing enterocytes at the tip of both villi (asterisks), and the concentric folds indicating contraction of the villus core and bunching of the surface epithelium in the neonate (representative of n=3, 1,000x, scale bar 30μm) (B) Neonate and juvenile villi following 30-minutes ischemia and 30-minutes *ex vivo* recovery. Note the sphering and persistence of microvilli (white arrowheads) in the neonatal wound-adjacent cells (left) versus the smoothened leading edges of the lamellipodia (white arrowheads) extending into the remaining defect (asterisks, wound edge outlined by white dotted line) in the juveniles (right). (representative of n=3, 5,000x and 10,000x, scale bars 10μm and 3μm)

### Neither a PGE_2_ treatment, the presence of the subepithelial compartment during *ex vivo* recovery nor *in vivo* recovery rescue neonatal restitution

Given the role of prostaglandins in driving villus contraction and tight junction restoration during subacute repair, we wondered whether the exogenous addition of prostaglandins during *ex vivo* recovery could support the restitution phase as well (1). To test this, 16,16 dimethyl-prostaglandin E_2_ was added to the Ringer’s solution in the basolateral reservoir bathing recovering neonatal tissue. Despite the addition of exogenous prostaglandin E_2_, there was a persistent 32±13.2% epithelial defect (Fig. 5 A, B). This resulted in no change in TEER (data not shown; P>0.05 for effect of recovery on TEER by two-way ANOVA) and no change in ^3^H-mannitol flux from the beginning to the end of the *ex vivo* recovery period (data not shown; P>0.05 by Sidak’s multiple comparisons test after one-way ANOVA). Additionally, endogenous production of prostaglandin E_2_ by recovering neonatal and juvenile mucosa during *ex vivo* recovery was measured and found to be similar across age groups (Fig. S1).

**Figure 5.**
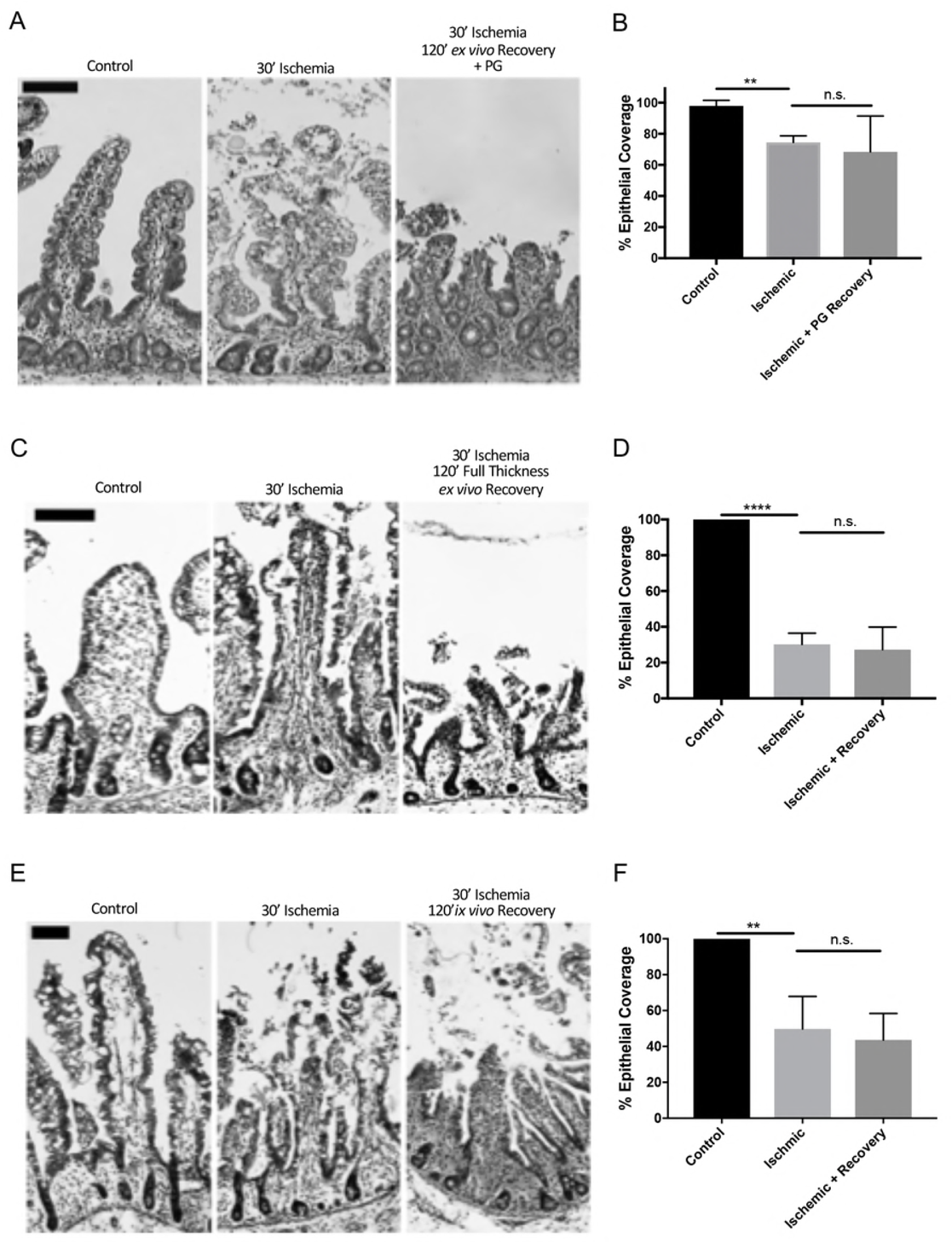
Effect of exogenous prostaglandins, full thickness *ex vivo* and *in vivo* recovery on neonatal restitution following 30-minutes of ischemia. (A) Representative histology of control, 30-minutes ischemic and 120-minutes *ex vivo* recovery neonatal jejunum with the addition of 1uM 16,16-dimethylprostaglandin E_2_ to the basolateral chamber. Note the persistent epithelial defect in the recovered tissue (scale bars 100 μm). (B) Histomorphometry quantified 74±2.5% and 68±13.3% epithelialization in injured and prostaglandin recovered tissues, respectively, as compared to 98±2.0% epithelialization of controls (n=3, n.s.= not significant, ^∗∗^P<0.01, unpaired t-test). (C) Representative histology of control, 30-minutes ischemic, and 30-minutes ischemic and 120-minutes full-thickness *ex vivo* recovery neonatal jejunum (scale bars 100 μm). (D) Histomorphometry quantified 30±6.3% and 27±12.6% epithelialization in injured and full thickness *ex vivo* recovered tissues, respectively, as compared to 100±0.0% epithelialization of controls (n=4, n.s.= not significant, ^∗∗∗∗^P<0.0001, unpaired t-test). (E) Representative histology of control, 30-minutes ischemic, and 30-minutes ischemic and 120-minutes *in vivo* recovery neonatal jejunum (scale bars 100μm). (F) Histomorphometry quantified 50±7.4% and 44±6.6% epithelialization in injured and *in vivo* recovered tissues, respectively, versus 100% epithelialization of controls (n=5-7, n.s.= not significant, ^∗∗^P<0.01, unpaired t-test).

To test whether the defect in epithelial restitution in neonatal tissues related to the lack of the seromuscular layers of the jejunum (and potential signals from these layers), neonatal experiments were repeated without separating the mucosa from the layers below prior to *ex vivo* recovery. Histomorphometry identified a 70±6.3% epithelial coverage defect from ischemic injury that persisted after full thickness *ex vivo* recovery (Fig. 5 C, D). This resulted in no change in TEER (data not shown; P>0.05 for effect of recovery on TEER by two-way ANOVA) and no change in ^3^H-mannitol flux from the beginning to the end of the *ex vivo* recovery period (data not shown; P>0.05 by Sidak’s multiple comparisons test after one-way ANOVA).

To further assess if the lack of recovery could be rescued with an intact mesenteric circulation, the ischemia/ recovery experiment was repeated with vascular clamps to reverse ischemia and reperfuse to recover the tissue *in vivo*. However, *in vivo* recovery resulted in similar TEER and ^3^H-mannitol flux in the ischemic and ischemic/ *in vivo* recovered tissues (data not shown, P>0.05 by one-way ANOVA for both analyses). Histomorphometry revealed a 50±7.4% and 56±6.6% epithelial coverage defect in ischemic and ischemic/ *in vivo* recovered tissues respectively (Fig. 5 E, F).

### Mucosal homogenate from ischemia-injured juvenile jejunum rescues neonatal repair

We hypothesized that the lack of recovery in neonatal tissues resulted from a lack of maturity of signaling elements within the mucosal compartment. To test this hypothesis, injured neonatal mucosal was recovered *ex vivo* in the presence of homogenized mucosal scrapings from 30-minute ischemia-injured juvenile versus neonatal jejunum. As expected, exogenous application of homogenized mucosa from injured neonatal jejunum failed to induce repair as per TEER measurements (Fig. 6A). However, application of homogenized mucosa from injured juvenile jejunum to both sides of the recovering neonatal tissue induced a robust increase in TEER in injured neonatal mucosa (Fig. 6A). Interestingly, application to either the basolateral or the apical side of the tissue did not induce any changes in the TEER (Fig. 6A). We then examined histological specimens prior to and following recovery to see if changes in TEER were associated with changes in epithelial coverage. This revealed an increase in epithelial restitution in injured neonatal tissues recovered in the presence of injured juvenile mucosal homogenate when applied both apically and basolaterally (Fig. 6 B, C). In contrast, application to either the basolateral or the apical side of the tissue, or the application of homogenized mucosa from injured neonatal jejunum did not produce the same effect. Histomorphometry indicated that tissues restituted to 80±4.4% epithelial coverage when exposed on both sides to the juvenile homogenate as compared to 40-60% in all other groups. Finally, to test whether the components within the injured juvenile homogenate responsible for inducing repair were soluble factors, the mucosal homogenates were centrifuged at 5,400 RCF and only the supernatant was applied to both sides of recovering neonatal mucosa *ex vivo*. Application of neonatal or juvenile supernatants produced no effect on TEER during 120-minutes *ex vivo* recovery (Fig. 7A). This was associated with no difference in epithelial coverage versus untreated tissue; there were persistent defects in restitution in all groups (Fig. 7 B, C).

**Figure 6.**
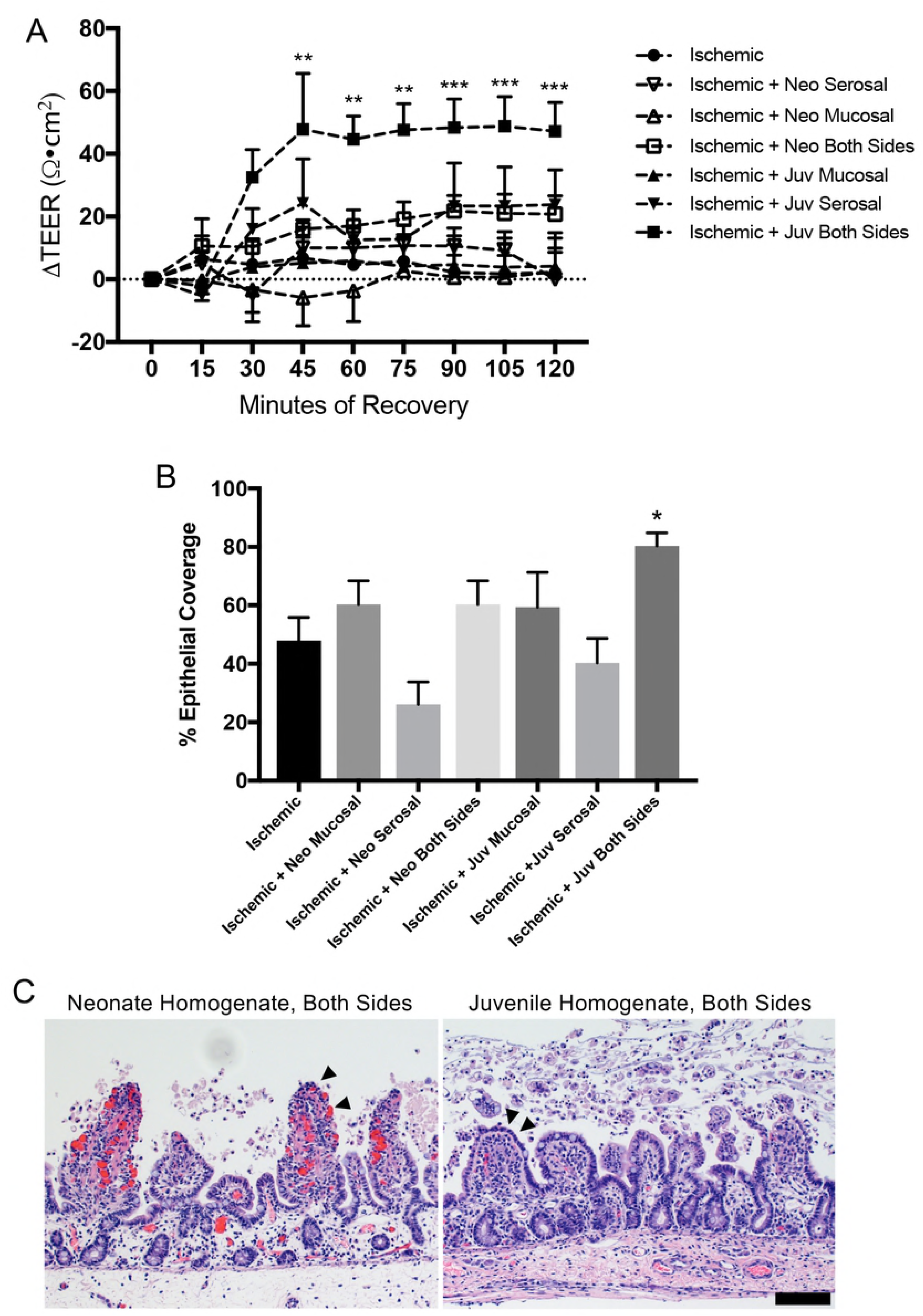
Exogenous application of injured juvenile mucosal homogenate partially rescues barrier repair in injured neonate jejunum. (A) Application of juvenile (juv), but not neonatal (neo), injured mucosal homogenate to both sides of the tissue during *ex vivo* recovery rescues the TEER of ischemia-injured neonatal jejunum. Application to apical or basolateral side only does not rescue TEER. Data presented is normalized relative to each individual tissue’s own initial TEER (n=5-6, significant effect of treatment and recovery on TEER by twoway ANOVA, P≤0.001; ^∗∗^P<0.01, ^∗∗∗^P<0.001 by Dunnett’s multiple comparisons test). (B) Representative histology shows remaining defects in neonatal homogenate-treated tissues as compared to evidence of partially restituted epithelium (arrowheads) in juvenile homogenate-treated tissues (n=6-12, scale bar 100μm). (C) Histomorphometry quantified 80±4.4% epithelial coverage with injured juvenile mucosal homogenate on both sides of the tissue versus 40-60% in all other treatment groups (n=6-12, significant effect of treatment on epithelial coverage by one-way ANOVA, P<0.01, ^∗^P<0.05 by Dunnett’s multiple comparisons test).

**Figure 7.**
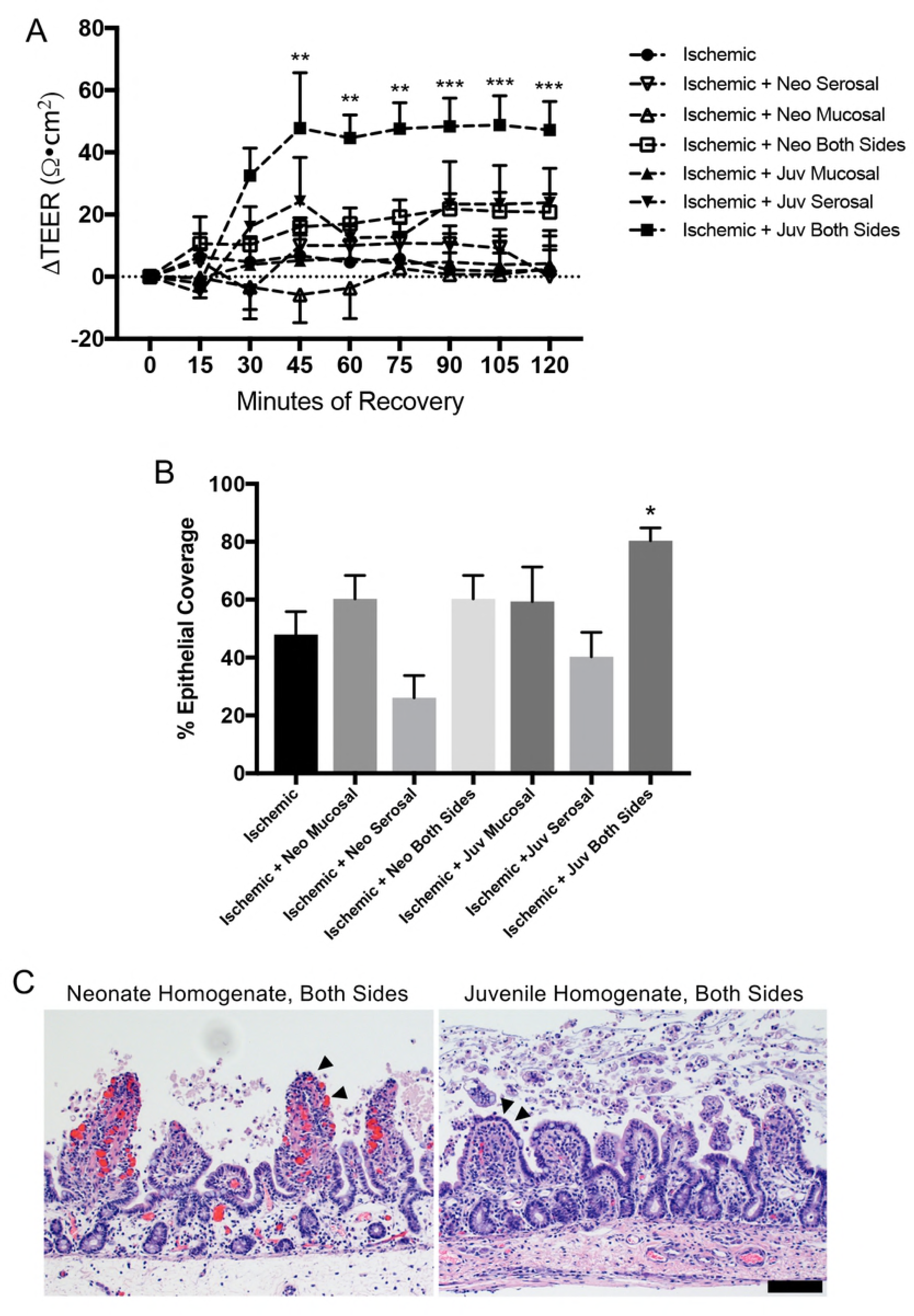
Exogenous application of neonatal or juvenile mucosal homogenate supernatant does not rescue barrier repair in injured neonatal jejunum. (A) Application of neither juvenile (juv) nor neonatal (neo) injured mucosal homogenate supernatant to both sides of the tissue during *ex vivo* recovery can rescue the TEER of ischemia-injured neonatal jejunum. Data presented is normalized relative to each individual tissue’s own initial TEER (n=5; significant effect of treatment but not recovery on TEER by two-way ANOVA, P<0.001; no significant differences versus Ringer’s control by Dunnett’s multiple comparisons test). (B) Representative histology shows remaining defects in neonatal homogenate-treated tissues as compared to evidence of partially restituted epithelium (arrowheads) in juvenile homogenate-treated tissues (n=6-12; scale bar 100μm). (C) Histomorphometry quantified 70±8.4% epithelial coverage in Ringer’s recovered tissue which did not differ from epithelial coverage in supernatant treated groups (61±11.0% and 71±11.3% for neonate and juvenile, respectively) (n=5-6; no differences by one-way ANOVA).

## DISCUSSION

NEC, volvulus and SIP are associated with mucosal disruption and inflammation due in part to intestinal ischemia (2). NEC is associated with estimated mortality rates between 20 and 30%, with the highest rates in neonates requiring surgery to resect intestine that fails to recover from injury (3, 4). In the subacute repair phase, epithelium adjacent to areas of wounding has been shown in animal models to efficiently restore barrier function by way of villus contraction, epithelial restitution, and tight junction restoration to limit sepsis, preserve intestinal viability and reduce host morbidity and mortality (1). However, in the present study, we have shown that there is a marked defect in the degree to which epithelial cells in neonatal pigs are able to restitute, and this is interestingly not the case in sow-matched piglets 6-weeks-of-age, which are only 4-weeks older. For example, quantifying restitution by histomorphometry following 30-minutes of ischemia in the two age groups revealed a stark difference in the percent wound closure between neonates (-11%) and juveniles (93%). Furthermore, juvenile animals demonstrate complete epithelial restitution and barrier restoration (as measured by TEER and ^3^H-mannitol flux) in tissues injured with up to 60-minutes ischemia and do this in a remarkably short period of *ex vivo* recovery (120-minutes). This complete barrier restoration following ischemic injury in our juvenile pig model is consistent with similar studies in the small intestine of adult human patients (19, 21). The disparity in repair between neonates and juvenile animals may partly explain high mortality rates associated with intestinal ischemic disease in infants.

One possibility that we considered to explain poor reparative responses in neonates in this study is that the neonatal mucosa may be time-dependently more susceptible to ischemic damage. One reason to consider this is the elongated height of villi in neonates, which would theoretically increase the counter-current exchange of oxygen within the villus vasculature that exacerbates ischemic injury at the tips of the villi(1,22). However, we were able to show that intestinal ischemia induces a similar degree of injury to the epithelium of both neonates and juveniles, relative to total villus length, as per our data demonstrating no significant difference in ischemia-induced loss of epithelial coverage in juvenile versus neonatal tissues. Furthermore, epithelial sloughing was shown to be restricted to the villus tips and the affected cells are mainly villin- and iFABP-expressing cells, indicating the injured cell population in both neonates and juvenile animals is composed of mostly differentiated enterocytes (20, 23). An important aspect of repair is villus contraction, which we thought might pose a significant challenge in neonates with such longer villi. However, neonatal villi undergo substantial contraction during recovery as evidenced by marked shortening of villus length by histomorphometry, creasing of the remaining epithelium as the villus core contracts on SEM, and an intact basement membrane on immunohistochemistry indicating there is no breakage of the subepithelial villus core.

The basement membrane was of added interest to us because of the critical role this plays in signaling and facilitating cell crawling. We examined the basement membrane of the wounded villi by labeling with collagen IV and appeared to be continuous and intact in both age groups. A defect in the basement membrane is thus likely not responsible for the impaired epithelial crawling noted in neonates. However, we did not examine any differences in the composition of other basement membrane constituents (nidogen, sulfated proteoglycans, and laminins) between neonates and juveniles. Notably, laminin isoforms, which are differentially expressed and often developmentally regulated, have differing regulatory properties for cell adhesion and migration and may be of interest for future study (24, 25).

Focusing specifically on epithelial crawling during restitution, enterocytes bordering the wound bed must depolarize and form lamellipodia to spread into and migrate across the defect until they contact other migrating epithelial cells, effectively closing the defect(1). We visualized this phenotype in recovering juvenile mucosa by SEM. More specifically, juvenile enterocytes at the wound margins could be seen assuming a migratory phenotype, flattening and spreading the redundant membrane of their microvilli into lamellipodia into the wound bed to close the defect. This was in marked contrast to the wound-adjacent enterocytes of neonates, which remained tall and round with intact microvilli, showing no evidence of spreading and lamellipodia formation. In addition, these cells did not appear columnar either, but rather appeared to have taken on a spherical brush-border-covered ‘tennis ball’-like appearance that suggested some level of cellular dysfunction. These findings suggest that neonatal enterocytes lack a mechanism of epithelial migration which may include the signals from the extracellular microenvironment, the appropriate receptors signal transduction pathways within the epithelium, the cellular machinery required for depolarization and directional migration, or the physical microenvironment required for adhesion and migration such as extracellular matrix or mucous layer components.

Previous work by our lab has implicated prostaglandin E_2_ in signaling for barrier repair in the context of first and last phases of subacute repair: villus contraction and tight junction reassembly(1, 26, 27). We therefore wondered if prostaglandin E_2_ also played a role in inducing epithelial restitution. However, ischemia-injured neonatal tissue treated with exogenous 16,16-dimethylprostaglandin E_2_ did not show improved recovery, and when this eicosanoid was measured in the apical Ringer’s solution sampled during *ex vivo* recovery, the levels detected did not differ between neonatal and juvenile pigs. Therefore, prostaglandin E_2_ does not appear to be a key missing factor responsible for the restitution defect in neonates.

This repair model employs *ex vivo* recovery and stripping of the seromuscular layers from the mucosa before mounting on the Ussing chambers, necessitating that the mucosa recovers independently of host input from any tissue beyond the muscularis mucosa. While this is not an impediment for the juvenile-aged animals, we wondered whether the addition of these host signals may rescue the failed restitution in the neonates. Therefore, we repeated the experiments in neonates without stripping the seromuscular layers for *ex vivo* recovery, reasoning that although this maneuver is intended to reduce the thickness of tissue which must be oxygenated and supplied with glucose *ex vivo*, neonatal tissues are thinner, and may be more likely to repair under full thickness conditions. However, there was no enhancement of the reparative response noted. In additional experiments, we took this idea a step further by reperfusing neonatal tissue for 120-minutes *in vivo* in order to allow potential factors from the systemic circulation and innervation to aid recovery, and to leave the tissue that much more intact to enable more physiological responses to injury. However, this experimental design also did not result in any notable improvement in mucosal repair and recovery of barrier function. Collectively, these data suggest that there is not a component of repair signaling deep to the muscularis mucosa, such as the host immune, nervous, or circulatory systems, that can rescue subacute repair in injured mucosal epithelium within neonatal animals. Importantly, these findings also confirm that the lack of recovery in neonatal mucosa *ex vivo* is not an artifact of the experimental conditions. Indeed, our model mirrors *in vivo* pathophysiology and is therefore a very powerful tool to study the development of neonatal intestinal repair mechanisms.

In light of systemic neonatal host inputs failing to rescue neonatal repair, we reasoned that the mucosal microenvironment from animals of the older age group might contain the elements required to promote repair. Indeed, this appeared to be the case because when exogenous homogenized mucosal scrapings from ischemia-injured jejunum from juveniles was placed in the Ussing chambers with ischemic-injured neonatal tissues, there was a robust recovery of TEER and a measurable increase in epithelial restitution in the neonatal mucosa. Importantly, this was not the case following application of homogenized mucosa from ischemia-injured neonatal jejunum, further supporting the presence of age-dependent pro-reparative factors within the juvenile milieu. These mucosal mixtures contain briefly homogenized mucosal scrapings consisting of everything from the level of the muscularis mucosa to the luminal surface including the live cells, cell secretory products, extracellular matrix and, very likely, the adherent mucous layer and microbiota. An interesting finding was that the homogenate had to be placed on both sides of recovering tissue to induce the restitution response. This may be due to a dose-response effect and application on both sides provides twice the dose of the key pro-repair factors. Alternatively, it may be that the key factors must act on both the basolateral and apical sides to induce a response, perhaps by acting on specific membrane-bound receptors on the epithelial cell, for example. Another interesting factor was that a low speed centrifugation to remove all solids from the juvenile homogenate attenuated the reparative effect of the homogenized mucosa, suggesting that the responsible components of the mixture either exist within the solids or are a labile secretory factor needing to be continually produced by live cells within the Ussing chamber reservoirs.

Future studies to identify the key factors responsible for producing the rescue effect in these experiments will further refine our mucosal repair model to encompass the postnatal development of restitution mechanisms. Paracellular signals from the lamina propria are known to modulate epithelial cell functions in homeostasis and disease states and cell populations within the subepithelial microenvironment are known to play a role in epithelial barrier repair signaling. For example, a growing body of evidence indicates that enteric glial cells (EGC) play a pivotal role in promoting intestinal epithelial repair and barrier function through the release of paracrine factors such as glial-derived neurotrophic factor, pro-epidermal growth factor, 11β prostaglandin F_2α_, and S-nitrosoglutathione (28–31). These EGC form a dense network in the lamina propria in close proximity to intestinal epithelial cells, and of particular interest, the EGC network of rodents has been shown to continue development after birth, with final maturation completing during the postnatal period (32–34). These features may implicate the EGC network in this neonatal repair defect. Key future experiments will examine the presence and role of glial-derived signaling in the defects of barrier repair in neonates. A differential transcriptome analysis of the juvenile and neonatal mucosal homogenate before and after ischemic injury can be expected to provide valuable insight to guide future work toward key signaling mechanisms and should be the next step in this line of work.

In conclusion, these studies show that there is a critical, but rescuable, defect in epithelial restitution following ischemic injury due to age-related insufficiency of repair mechanisms within the mucosal compartment. These findings may be of significant translational relevance to high morbidity and mortality in pre-term and term infants afflicted with ischemia-related intestinal disease. This defect may also be relevant to other mechanisms of intestinal epithelial injury, expanding its applicability in numerous other intestinal diseases of neonates. Identifying the specific rescuable defects in repair mechanisms will enhance our understanding of the development of barrier repair mechanisms in the postnatal period and will guide future clinical interventions to improve outcomes in affected infants.

## ACKNOWLEDGEMENTS

The authors want to thank Gwendolyn Carnighan, Erik Wegner-Clemens, Megan Gallagher for their assistance with experimental procedures and data analysis, the staff and faculty of Lab Animal Resources and the Central Procedures Laboratory at NC State College of Veterinary Medicine for their excellent animal care and assistance with surgical procedures, the NC State Swine Educational Unit for their logistical support and exceptional animal care, and Victoria Madden and Kristen White for providing their expertise in electron microscopy.

## AUTHOR CONTRIBUTIONS

Conceived and designed all experiments: ALZ, ATB, LVL, JO. Performed the experiments: ALZ, TAP, JKM, LMG, ATB. Contributed materials, equipment and analysis tools: JO, LMG, LVL, ATB. Prepared the final manuscript: ALZ, ATB.

**Supplemental Figure 1.**
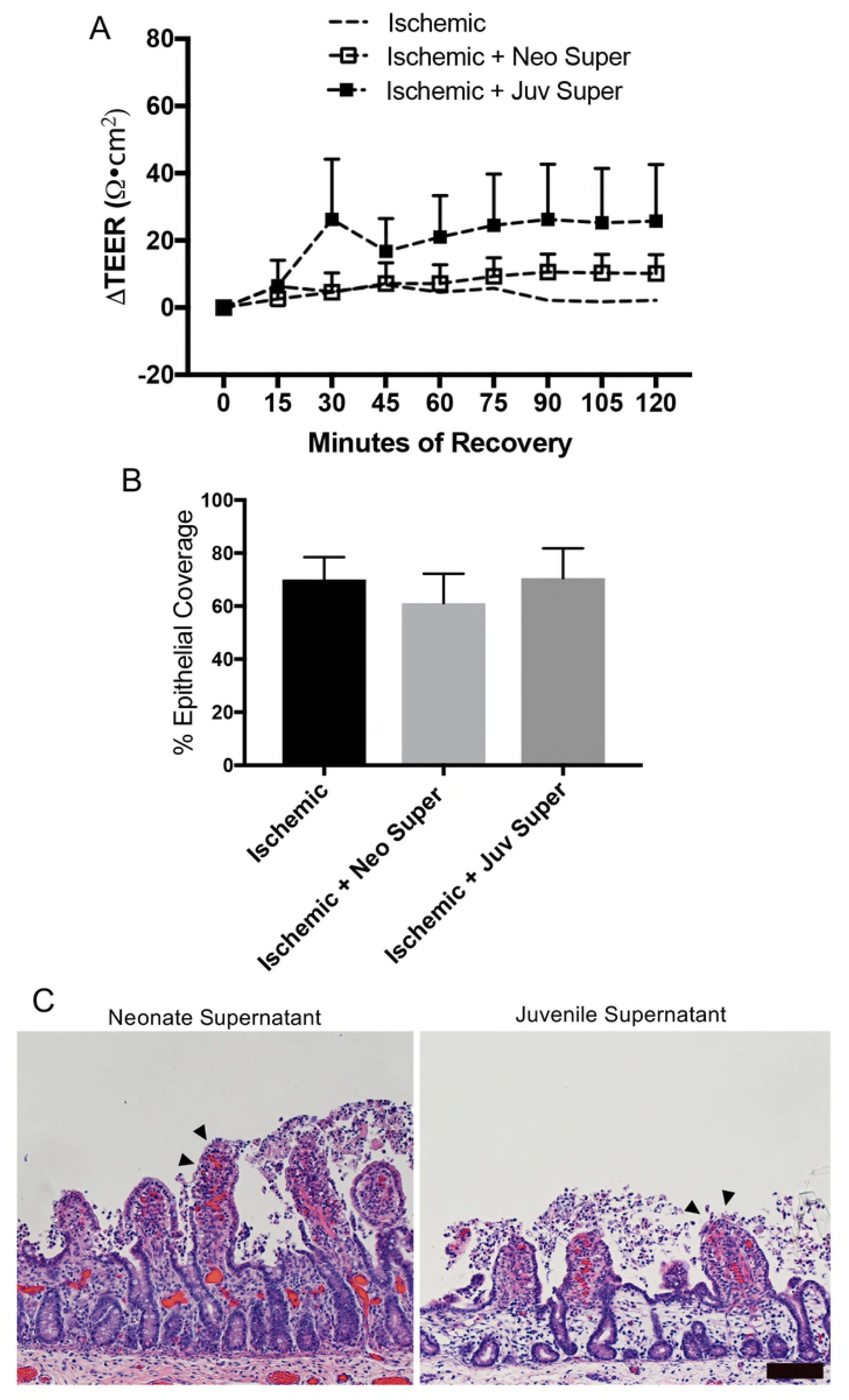
Injury induced similar PGE_2_ production in the jejunum of neonates and juveniles. PGE_2_ production in the basolateral Ringer’s solution at 60-minutes of *ex vivo* recovery was induced by ischemic injury similarly across both age groups. (n=5; P<0.001 for effect of injury on PGE_2_ by two-way ANOVA; n.s. = no significant difference on Sidak’s multiple comparisons test).

